# Coexistence patterns and diversity in a trait-based metacommunity on an environmental gradient

**DOI:** 10.1101/2021.06.15.448245

**Authors:** Mozzamil Mohammed, Bernd Blasius, Alexey Ryabov

## Abstract

The dynamics of trait-based metacommunities have attracted much attention, but not much is known about how dispersal and spatial environmental variability mutually interact with each other to drive coexistence patterns and diversity. Here, we present a spatially-explicit model of resource competition in a metacommunity on a one-dimensional environmental gradient. We find that both the strength of dispersal and the range of spatial environmental variability affect coexistence patterns, spatial structure, trait distribution and local and regional diversity. Without dispersal, species are sorted according to their optimal growth conditions on the gradient. With the onset of dispersal source-sink effects are initiated, which increases the effects of environmental filtering and interspecific competition and generates trait lumping, so that only a few species from an environment-defined trait range can survive. Interestingly, for very large dispersal rates the system becomes spatially homogeneous, but nevertheless two species at the extreme ends of the trait-off curve can coexist for large environmental variability. Local species richness follows a classic hump-shaped dependence on dispersal rate, while local and regional diversity exhibit a pronounced peak for intermediate values of the environmental variability. Our findings provide important insights into the factors that shape the structure of trait-based metacommunities.

## Introduction

Ecological theory has advanced our understanding of how diversity patterns are shaped by the joint influence of coexistence mechanisms and spatial structure. In particular, spatial environmental variability has long been considered as an important coexistence mechanism by which ecological communities are spatially structured and diversity patterns are generated. In this context, the paradigm of a metacommunity, defined as a set of local communities connected by dispersal (Leibold et al. 2004), has evolved into one of the most successful approaches for describing spatial ecosystems, allowing to link local coexistence theory and spatial processes.

According to the metacommunity paradigm, patterns of species coexistence and diversity are governed by four major archetypes: species sorting, patch dynamics, mass effects and neutral dynamics (Leibold et al. 2004). These mechanisms rarely act in isolation and some advances have been made towards synthesizing their effects (Leibold 2011, Fournier et al. 2017, Thomson et al. 2020, Bauer et al. 2021). A recently proposed metacommunity framework (Thomson et al. 2020) reduced the number of factors that are shaping ecological communities to the interplay of three fundamental processes: (i) Density-independent growth rates that are based on abiotic conditions and vary in space, (ii) density-dependent biotic interactions, and (iii) dispersal. Thereby, dispersal generates source-sink effects: emigration locally reduces population size in sites where conditions are more favorable, whereas immigration increases population size in less favorable sites, and introduces species that interact with local biota, thus altering density-dependent processes and competition.

Most modelling studies of competitive metacommunities focus on trait-based approaches where each species is characterized by the mean value of its functional traits (McGill et al 2006, Litchman and Klausmeier 2008, Bauer et al. 2021). One popular conceptual model is that of competition for two limiting resources (Leon & Tumpson 1975, Grover 1997, Tilman 1982, Chase and Leibold 2009). Here, the species’ traits determine species preference for one of the two resources, the density-independent abiotic conditions (sensu Thomson et al. 2020) are determined by the supply and ratio of resources, and density-dependent species interactions follow from consumption and competition for resources. In the simplest case of only one limiting resource, the species with the highest competitive ability for that resource will outcompete all other species, while in a two species system with two limiting resources coexistence and priority effects (bistability) are possible (Leon and Tumpson 1975). The competition outcome can be reverted in a spatially extended system, where parameter combinations that lead to coexistence in a uniform environment may favor alternative stable states in a spatial system, and vice versa (Ryabov and Blasius 2011, Tsakalakis et al 2020). Extensions of resource competition models for many consumers typically assume that species traits are distributed along a trade-off curve. Then, a typical competition outcome is the survival of a single species whose traits are best matched to the ratio of resources supplied (Koffel 2016).

Resource competition models have frequently been applied to describe metacommunities (Abrams and Wilson 2004, Mouquet et al. 2006, Hodapp et al. 2016, Wickmann et al. 2017, Wickman et al. 2019). In such spatial settings, dispersal becomes an important additional factor for controlling diversity and community structure. While some studies also consider for flow of resources between local sites, going over to a meta-ecosystem (Haegeman and Loreau 2014, Tsakalakis et al 2020), most modelling studies of resource competition in metacommunities restrict dispersal to emigration and immigration of consumers. It has been suggested that in such systems dispersal profoundly affects local and regional diversity patterns and that the effect depends on the degree of spatial environmental heterogeneity (Kneitel and Miller 2003, Mouquet et al. 2006, Haegeman and Loreau 2014).

Dispersal strength depends on species dispersal traits and on the spatial arrangement of habitat patches in the landscape. Dispersal can move individuals from suitable into unsuitable habitats where their expected fitness becomes low, potentially causing local extinctions and decreasing the total biomass. One the other hand, dispersal may also increase fitness by sending species into more suitable habitats where traits of species can match the environmental conditions, or the interspecific competition is weak (Ryabov and Blasius 2008). Dispersal-driven environmental filtering can thus alter the total biomass and the spatial distributions of species. For small dispersal, the bulk of each species biomass will be concentrated around optimal growth conditions and the species will be sorted according to their traits along a spatial gradient. Increasing the strength of dispersal generates source-sink effects, promoting spatial homogenization (Mouquet and Loreau 2003). Thereby, for intermediate dispersal rates the local diversity enhancing effect of immigration dominates (as new species arrive in each locale), while in the limit of large dispersal rate local extinctions occur due to the higher interspecific competition and environmental filtering. In total, these effects yield a hump-shaped dependence of local diversity on dispersal, characterized by highest local diversity for intermediate levels of dispersal (Mouquet and Loreau 2002, 2003).

Similar effects arise for gradients of environmental heterogeneity, where highest diversity is obtained for intermediate levels of environmental variability (Kunin 1998, Mouquet and Loreau 2002, Mouquet et al. 2006). Despite this growing recognition about the role of dispersal and environmental variability in isolation, their joint influence has rarely been studied (Mouquet et al. 2006, Tsakalakis et al. 2020) and a general understanding of how regional heterogeneity and dispersal interact is still missing.

This lack of knowledge is partly due to the overwhelming complexity of possible topological arrangements of patch networks and the corresponding distribution of environmental conditions in the landscape (Fournier et al. 2017) which poses major technical difficulties for a comprehensive theoretical investigation. One convenient way to reduce this complexity is to restrict the analysis from general habitat networks to spatial gradients, in which an environmental state varies continuously along a one-dimensional spatial coordinate. The theoretical description of metacommunities on a spatial gradient is rooted in models from quantitative genetics (Kirckpatrick and Barton 1997; Case and Taper 2000), describing the spatial range of a species with a certain quantitative character. Subsequent studies (Doebeli and Dieckmann 2003, Leimar et al. 2008, Norberg et al. 2012) revealed the ubiquitous emergence of trait lumping on the spatial gradient – a finding that contradicts the intuitive perception that gradual spatial variation in environmental conditions should result in a gradual variation of trait values. Similar to spatially implicit models on a single trait-axis (Scheffer and van Nes 2006) the emergence of such trait clusters in trait-space dimensions can be understood with approaches from pattern formation theory (Pigolotti et al. 2007, Delfau et al. 2016).

A recent modelling study investigated resource-based competition of a metacommunity on a spatial environmental gradient (Hodapp et al. 2016). The study revealed a strong dependence of diversity patterns on spatial resource variability, in particular, on the match of environmental and trait variability. These results are in line with the intuitive expectation that a strong positive relationship between diversity and ecological function (here, resource use) requires that a large environmental variability is met by a large variation in traits (Ptacnik et al. 2010). While Hodapp et al. (2016) studied the relation between trait and environmental variability, few studies investigated how this relation is influenced by varying dispersal strength. A first study was performed by Mouquet et al. (2006), who found that the highest diversity occurs at intermediate levels of dispersal; but this effect was dependent on the environmental heterogeneity, with highest values of regional and local species richness for intermediate levels of environmental heterogeneity. This study was restricted to two local patches and a small species pool and thus was not able to resolve spatial structure or to draw conclusion about the possibility of niche lumping. Thus, many questions regarding the relationship of dispersal and environmental variability in resource-based metacommunities along an environmental gradient, so far, remain unanswered.

In this study, we develop and analyse a spatially-explicit resource-competition model of competing populations on an environmental gradient. Our model describes the situation of randomly dispersing consumers that compete for two heterogeneously distributed resources (Hodapp et al 2016). We analyse the joint effect of dispersal and variability in spatial resource supply (the steepness of the environmental gradient) on coexistence patterns, species sorting, trait-distributions, as well as local and regional diversity.

We find that without dispersal, species are sorted according to their optimal growth condition on the gradient, but with the onset of dispersal source-sink effects are initiated. Thereby, the dispersal rate and the range of spatial environmental variability strongly affect the competition outcomes, composition, and diversity. That is, at low dispersal rates the number of surviving species increases with the spatial environmental variability. Increasing dispersal rates generates trait lumping and strengthens the effect of environmental filtering and interspecific competition so that only a few dominant species can survive. Interestingly, for very large dispersal rates the system becomes spatially homogeneous, but nevertheless two species at the extreme ends of the trait-off curve can coexist. Our model also provides important insights into the factors that shape metacommunity structure and promote coexistence. According to our simulations global species richness depends in an intricate manner on dispersal strength and resource variability, with a classic hump-shaped dependence of diversity on dispersal rate, but also a pronounced peak of global diversity for intermediate values of resource variability.

## Model and method

### Model description

We use a spatially-explicit resource-competition model, based on the model framework described by (Mouquet et al. 2006, Hodapp et al. 2016). We assume that species live in a one-dimensional spatial environment that is discretized into *k* independent patches of equal sizes. Each patch contains two essential resources R_*j*_(j = 1,2) supplied locally and shared by a community of *n* species. We assume that the resource supply ratio changes from left to right along the environmental gradient (Fig. 1). Species move between adjacent patches with dispersal rate *d*, and compete for the resources in each patch. Let N_*i,l*_ denote the local biomass of species *i* = 1 ... *n* and R_*j,l*_ the local concentration of resource *j* = 1,2 in patch *l* = 1 ... *k*. The growth g_*i,l*_ of species *i* in patch *l* is determined by the availability of resource 1 and 2, and follows a Monod kinetics with Liebig’s law of the minimum (León & Tumpson 1975, Grover 1997, Tilman 1982, Chase and Leibold 2009)

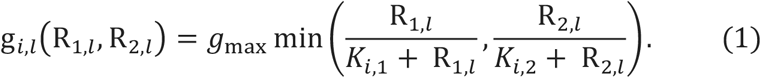

**Fig. 1:**
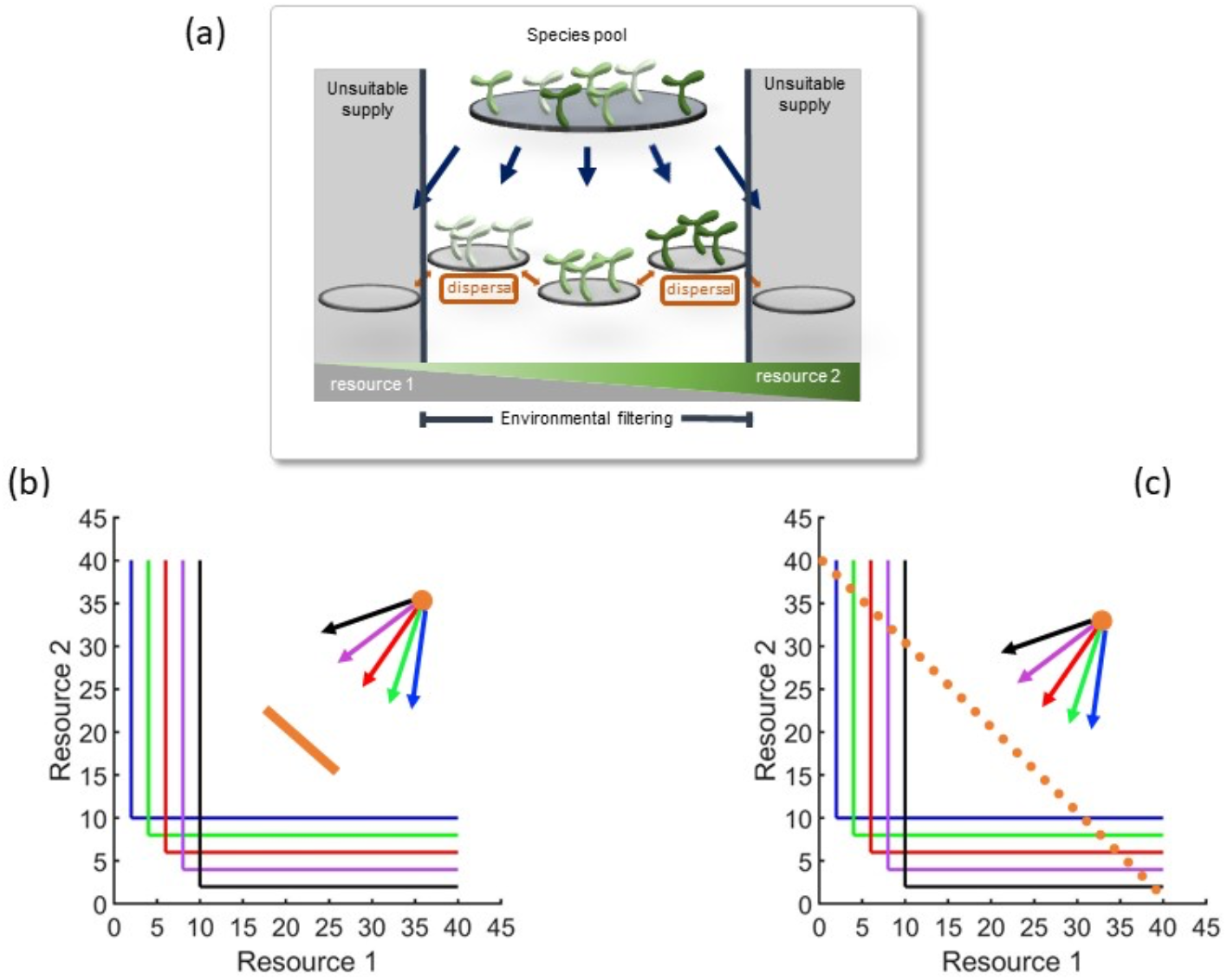
(a) Conceptual framework depicting the metacommunity response to spatial resource distributions, dispersal and environmental filtering. Consumers live, and can locally disperse, on a chain of discrete patches which are provided a varying supply of resources 1 and 2 from left to right. Growth rates are determined by the match of species’ resource requirements and local resource availability, which can yield unsuitable habitats at the ends of the gradient (environmental filtering). (b, c) Competitor trade-off in resource requirements. Shown are the zero net-growth isoclines (perpendicular solid and dashed lines) and the direction of consumption vectors (top right) for five species (different colors) in the resource plane. Orange circles indicate the resource levels, (*S*_1,*l*_, *S*_2,*l*_), that are supplied at different spatial positions. The scatter range of these supply points expresses the degree of environmental variability. In panel (b) the spatial environmental variability is low (orange circles are close to each other), while in panel (c) the spatial environmental variability is large.

Here *g*_max_ is the maximum reproduction rate and *K*_*i*,1_ and *K*_*i*,2_ are the half-saturation constants of species *i* for resource 1 and 2, respectively.

In the absence of dispersal (*d* = 0) the net growth rate of each species is given by the difference between reproduction and mortality rates. The balance between reproduction and mortality, g_*i,l*_(R_1,*l*_,R_2,*l*_) = *m*, defines the so-called Zero Net Growth Isoclines (ZNGIs) which divide the resource plane for each species *i* into areas of positive and negative growth (Grover 1997, Ryabov and Blasius 2011). Solving this equation we obtain the minimal resource requirements of species *i* for resource *j* (the *R*\*values, Tilman 1980)

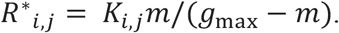

To generate trait variability, we parameterize the species’ half saturation constants in such a way that the respective *R*\*values are equally distributed on a one-dimensional straight trade-off line in the resource plane (Figs. 1b and c). Thereby, for simplicity, we keep *g*_max_ and *m* identical for all species and independent of the environment. We denote a species close to one of the edges of the trade-off line as *extreme*, as it has low requirements for one of the two resources. In contrast, we denote species with traits in the interior of the trade-off line as *moderate*.

To model metacommunity dynamics, we allow that individuals disperse between adjacent patches. The growth of species *i* in patch *l* then follows the equation

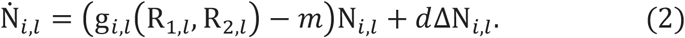

Here, the second term is the discrete Laplace operator which describes the diffusion of individuals of species *i* on a one-dimensional lattice with cell size *h* (here set to 1)

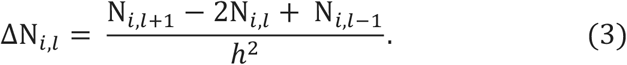

We assume that species do not disperse beyond the boundaries of the lattice (i.e., patches 1 and *k*). That is, species in patch 1 can only disperse to the right of its patch while species in patch *k* can only disperse to the left of its patch.

To generate an environmental gradient, we parameterize the external supply *S*_*j,l*_ of resource *j* in patch *l* in such a way that *S*_1,*l*_ linearly decreases and *S*_2,*l*_ linearly increases with the spatial patch position *l* (Fig. 1a). We denote the difference between the highest and the lowest concentration of resource supply on the gradient as the resource variability Δ*S*. The resource variability defines the steepness of the environmental gradient and is adjusted by varying the maximal range of resource supply points in the resource plane (orange circles in Figs. 1b and c).

We do not consider diffusion of resources between patches, and thus, the dynamics of resource concentrations are governed by the difference between resource inflow and consumption

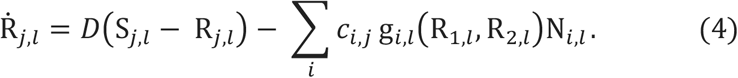

Here, *D* is the dilution rate and *c*_*i,j*_ the consumption rate of resource *j* by species *i*, indicating the resource amount required for a consumer to reproduce a unit of biomass. We define the consumption vector based on optimal foraging theory (Tilman 1982), which suggests that each species consumes a resource in proportion to its minimal requirements for this resource; thus *c*_*i,j*_ = *αR*\*_*i,j*_, where *α* is a constant and set to 0.05.

### Numerical scheme

The developed metacommunity model is given as a system of ordinary differential equations (Eqs. 1–4). To numerically solve this system, we used the ode45 function of Matlab 2019. We confirmed its robustness by using the Differential Equations package (Rackauckas and Nie, 2017) of the programing language Julia (Bezanson et al. 2017), yielding very similar results. As initial values we assumed that all species were present in each local site at an initial biomass randomly distributed in the range from 1 to 20, while the initial concentrations of resources ranged from 1 to 20.

To compute steady state solutions, we used an optimized numerical scheme. For this we first ran the simulation for a transient of 10,000 units of time to obtain a solution close to the equilibrium of the dynamical system. Next, we used this obtained solution as the initial guess for a root-solving algorithm (the Matlab function fsolve) to numerically compute the equilibrium solution of Eqs. (2) and (4).

### Diversity estimation

We computed the local and regional diversity of the simulated metacommunities based on the Shannon diversity index. The Shannon index for regional diversity was computed as *H*_reg_ = − ∑_*i*_ *p*_*i*_ln (*p*_*i*_), with *p*_*i*_ = *N*_*tot,i*_/∑_*i*_ *N*_*tot,i*_ the relative total biomass of species *i* over all patches and *N*_*tot,i*_ = ∑_*l*_ *N*_*i,l*_ the total biomass of species *i* over all patches. Computation of local diversity was based on the patch-specific relative biomass *q*_*i,l*_ = *N*_*i,l*_/∑_*i*_ *N*_*i,l*_. We computed the average relative biomass of species *i* over the *k* patches as *q*_*i*_ = ∑_*l*_ *q*_*i,l*_/*k*, yielding the Shannon index for local diversity as *H*_loc_ = − ∑_*i*_ *q*_*i*_ln (*q*_*i*_). Finally, we computed the effective species numbers as *E*_reg_ = exp(*H*_reg_) and *E*_loc_ = exp(*H*_loc_), representing the number of equally common species present in the environment (resp. regionally and locally) required to yield the same Shannon index (Jost 2006).

**Table 1:** Variables and parameters used in the model. Following Hodapp et al. (2016) we used dimensionless units for length, time, biomass and resource concentrations. Values for maximum growth, mortality, consumption and dilution rates were taken from Hodapp et al. (2016), but we increased the lattice size to *k* = 50 and restricted R* values in the range 2-10.

| Symbol | Description | Default value |
| --- | --- | --- |
| $N_{i,l}$ | Biomass of species $i$ in patch $l$ | |
| $R_{j,l}$ | Concentration of resource $j$ in patch $l$ | |
| $g_{i,l}$ | Growth rate of species $i$ in patch $l$ | |
| $g_{\text{max}}$ | Maximum growth rate | 1 |
| $m$ | Mortality rate | 0.25 |
| $R^*_{i,j}$ | Minimum resource requirement of species $i$ for resource $j$ | 2-10 |
| $K_{i,j}$ | Half-saturation constant of species $i$ on resource $j$ | $R^*_{i,j} \frac{g_{\max} - m}{m}$ |
| $c_{i,j}$ | Consumption rate of species $i$ on resource $j$ | $0.05 * R^*_{i,j}$ |
| $S_{j,l}$ | Supply of resource $j$ in patch $l$ | $\left[ \bar{S} - \frac{\Delta S}{2}, \bar{S} + \frac{\Delta S}{2} \right]$ |
| $\Delta S$ | Resource variability | 1- 40 |
| $\bar{S}$ | Mean resource variability | 20.5 |
| $D$ | Dilution rate of resource $j$ | 0.25 |
| $k$ | Number of patches | 50 |

## Results

### Effects of increasing dispersal rate

We begin our analysis by investigating how the competition outcome and equilibrium diversity of a metacommunity on an environmental gradient depend on the combined interplay of dispersal rate and resource variability. To obtain conceptual insights, we first restrict the analysis to a small-sized metacommunity of only three competitors (one ‘moderate’ and two ‘extreme’ species) and later extend the analysis to a larger number of species. Fig. 2 depicts our simulation results, where we model the metacommunity in a wide range of dispersal rates (over six orders of magnitude) and in a gradient from small to large resource variability. We find that at low dispersal rates all three competitors can coexist regionally, but the level of resource variability determines environmental filtering and affects species evenness. When resource variability is small, the moderate species is relatively more competitive and attains higher equilibrium biomass (Fig. 2 a). In contrast, the extreme species on average have smaller biomass, as they thrive in conditions of unbalanced resource supply which are not present in the lattice for low environmental variability. With increasing dispersal strength this unbalance is even enhanced by mass effects so that eventually the extreme species are outcompeted by the moderate species. This outcome is reversed by increasing resource variability (Figs. 2 b and c) because it creates local patches with extreme environmental conditions, favoring the extreme species. Accordingly, increasing the range of resource variability weakens the effect of environmental filtering on the extreme species and increases their biomass, to the extent that this time the moderate species (rather than the extreme species) is outcompeted at large dispersal rates (Figs. 2 c).

**Fig. 2:**
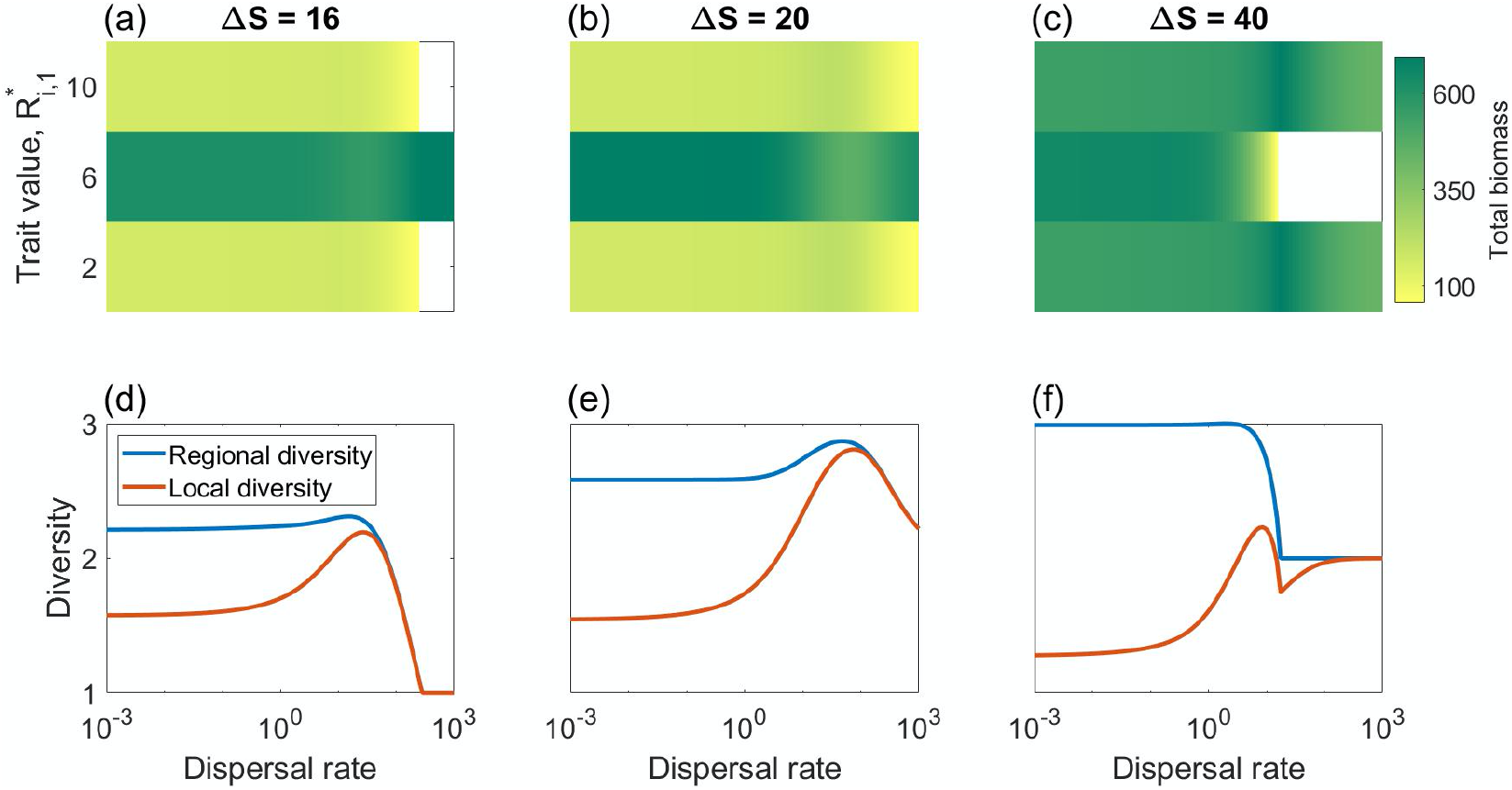
Competitive outcome and emerging diversity in a metacommunity of three consumers in response to spatial environmental variability ΔS and dispersal rate. (a-c) The total biomass *N*_*tot,i*_ of consumer *i* in equilibrium (color coding) and (d-f) the corresponding local and regional effective species number (red and blue lines) as a function of the dispersal rate for three levels of spatial resource variability (Δ*S* =16, 20 and 40). The y-axis in (a-c) represents the trait value of a consumer *i* as its requirement, *R*\*_*i*,1_, for resource 1. We use the values *R*\*_*i*,1_ = 2, 6, and 10, and *R*\*_*i*,2_ = 10, 6, 2, for species 1, 2 and 3, respectively. See Table 1 for other parameter values.

This shift in species dominance is also visible in the dispersal dependence of local and regional diversity (Fig. 2, bottom panels). When dispersal is small, local diversity *E*_loc_ ranges between 1 and 2 (red curve). This can be explained by the fact that when the patches are weakly coupled the species are spatially sorted along the environmental gradient and no more than two species coexist in each patch. With increasing dispersal, biomass flows from source to neighboring patches are initiated, increasing local diversity there due to the mass effect. Increasing dispersal even further, these locally co-occuring species start to outcompete each other, eventually reducing local diversity again – yielding, in total, in a hump-shaped relationship between local diversity and dispersal rate (Mouquet and Loreau, 2002). Unlike the local diversity, regional diversity *E*_reg_ attains large values at small dispersal rates when each species occupies its source patch (Fig. 2, bottom panels, blue curve). With increasing dispersal, the regional diversity at first slightly increases (reflecting the more even community composition), but then decreases (when outcompeted species are lost from the whole metacommunity) and merges with the local diversity at large dispersal rates (when the system becomes spatially homogenous so that differences between local and regional diversity are obliterated).

These results for a three-species metacommunity basically remain unchanged when we extend the species pool to a metacommunity of 15 species with the same range of trait variability (Fig. 3). At low dispersal rates, we again observe a low local diversity of about *E*_loc_ = 1, but for small resource variability moderate species are favored so that only species with trait values in the middle of the trade-off curve are filtered by the environment and survive (Fig 3a). With increasing resource variability, the system contains local patches with more and more unbalanced resource supply, favoring extreme species and increasing the number of surviving species and their range of traits (Figs. 3 b and c). With increasing dispersal rate, local diversity in first enhanced due to mass effects, while at the high end of dispersal local diversity decays again as an increasing number of species are outcompeted, yielding a hump-shaped local diversity curve (Fig. 3, bottom panels). In the same way, quite analogous to the three-species case, also in the 15-species metacommunity regional diversity starts out with a high value for small dispersal rate and decays to the base line of the local diversity for larger dispersal rates when the system becomes spatially homogeneous.

**Fig. 3:**
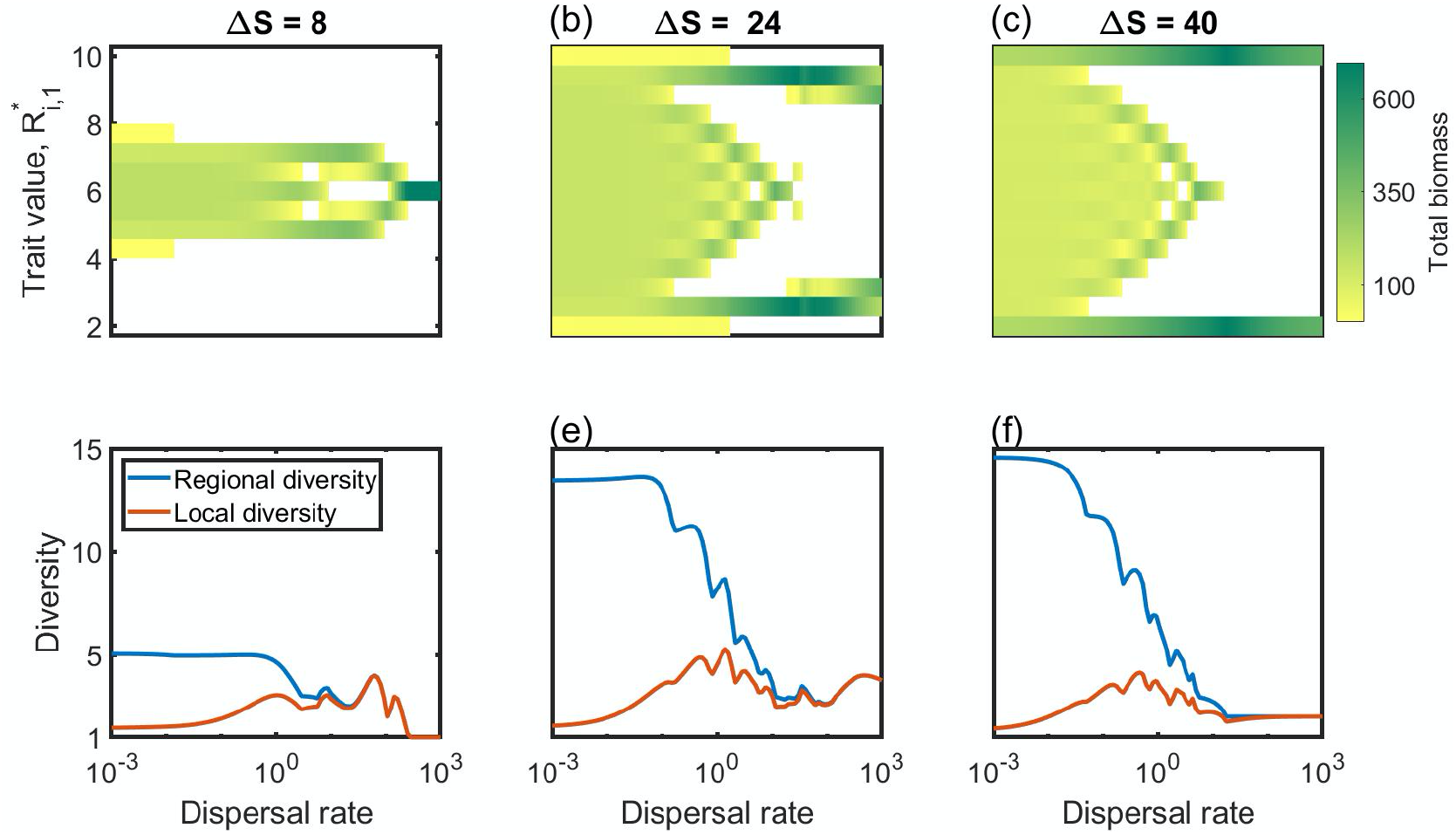
The same as described in the caption of Fig. 2, but here shown for a metacommunity of 15 species. Consumer resource requirements were equidistantly distributed in the range *R*\*_*i*,1_ = 2 . . 10 and *R*\*_*i*,2_ = 10 . . 2.

Despite these similarities, our simulations also reveal some marked differences in the competition outcome between the small and the large meta-community. The reason for this is that the model with 15 species allows us to resolve finer details in the composition of traits, revealing some intriguing patterns of how the trait distribution changes with dispersal rate. This becomes particularly apparent in the range of intermediate traits, where with increasing dispersal the range of the surviving traits is shrinking in a tongue-like pattern. The equilibrium trait distribution within this 'trait tongue’ is not homogeneously filled, but instead, we observe an alternating pattern of surviving traits, reminiscent to trait lumping (Scheffer and Nes, 2006, Leimar et al. 2006).

Beside this fine-structure in the emerging trait distributions, the 15 species metacommunity also exhibits some distinctive large-scale patterns (Fig. 3). Species with traits directly outside of the trait tongue do not survive, yielding some ‘forbidden’ zones in the trait-dispersal plane, visible as ‘white space’ patterns in Fig. 3. Only at the extreme ends of the trait range species can survive, but only if resource heterogeneity is sufficiently strong (Fig. 3 b and c). As a result, in the limit of large dispersal metacommunity structure seems to be restricted to two characteristic cases. The first case arises for small resource variability. Then, a single moderate species in the middle of the trait-tongue survives and dominates the community. If, however, resource variability is sufficiently large so that extreme species are able to survive, these will outcompete the moderate species. As a result, for intermediate or large resource variability, the two extreme species at the opposing ends of the trade-off line coexist and dominate the community.

### Biomass distributions in the space-trait plane

More insight into the origin of these patters is obtains by plotting the emerging equilibrium distributions of biomass in the space-trait plane. The case of a large species pool (15 species) with high resource variability (Δ*S* = 40), shown in Fig. 4 for different dispersal rates, gives the best insight into how patterns in physical space (spatial structure) and in trait-space (community composition) mutually depend on each other. At low dispersal rates, all species are linearly sorted along the gradient according to their optimal trait value at each location (Fig. 4a). With increasing dispersal, species are horizontally dispersed from their optimal patches to location of unfavorable conditions (mass effect), so that biomass spreads over a larger number of patches (Figs. 4b and c). As a result, species of different trait values appear on the same ‘vertical’ in this figure, i.e., they co-occur and compete locally. When this causes species to be locally suppressed, or even outcompeted, in a patch, small gaps in the trait-distribution appear. These gaps organize themselves over the whole trait distribution to form trait lumps (Scheffer and Nes, 2006, Leimar et al. 2006), as is apparent in Fig. 4c in the regular vertical spacing of high-biomass (green) stripes which are intersected by low-biomass (yellow) stripes. At the same time, with increase of dispersal rate, at the extreme ends of the trait range large gaps appear. This means that the most competitive species have traits either in the middle or at the edges of the trait range, while species with traits between the extreme and middle traits are outcompeted (Fig. 4c). Finally, at large dispersal rates, the two extreme species spread across the entire environment and outcompete all species with moderate traits (Fig. 4d). Note, the different competition outcome, if resource variability is smaller (Δ*S* = 8) as shown in SI Fig. 7. In this case, first the realized trait range is smaller than for the case with large resource variability (Fig. 4). But most importantly, for small resource variability in the limit of large dispersal the metacommunity is dominated by a single moderate species (Fig. 7d), whereas for large resource variability the metacommunity is dominated by a coexistence of two extreme species (Fig. 4d).

**Fig. 4:**
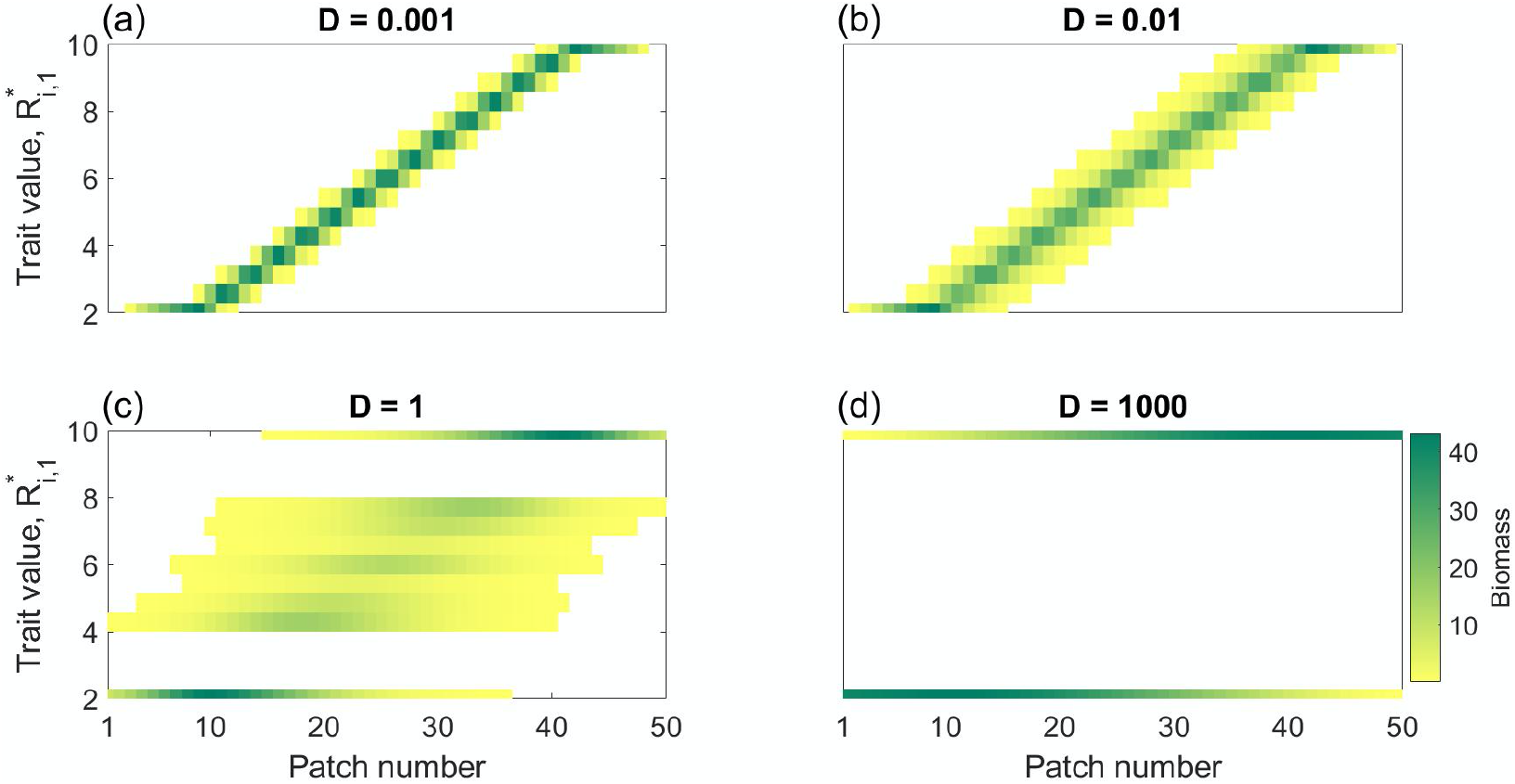
Influence of dispersal on equilibrium space-trait distribution of biomass. The figure shows the local biomass (color coding) of a metacommunity of 15 species in equilibrium in dependence of the spatial position on the gradient (patch number) and the trait value (requirement, *R*\*_*i*,1_, for resource 1) for four different levels of dispersal. (a) Low dispersal rate (*D* = 0.001), species are linearly sorted along the gradient to positions of optimal fitness. (b) Small dispersal rate (*D* = 0.01), spatial ranges of species start to extend in space. (c) Intermediate dispersal rate (*D* = 1), spatial ranges of a species extend over the whole domain, emergence of trait lumping. (d) Large dispersal rate (*D* = 1000), only the best competitors survive and spread over the whole environment. Parameter values as in Fig. 3c (Δ*S*=40).

### Influence of resource variability

To highlight effects of the dispersal-driven environmental filtering on the community composition, consider the equilibrium trait distributions of the large (15 species) metacommunity in dependence of the resource variability for four disparate levels of dispersal rates (Fig. 5). At low dispersal rate (Fig. 5 a and b) we observe a uniform regional distribution of trait values within a finite range. This trait range of surviving species becomes larger with increasing resource variability, as more and more local patches with extreme conditions of unbalanced resource supply are present in the system, allowing more extreme species to grow. In our parametrization, starting from a resource variability of about 20 to 30, this trait range covers the whole species pool, that is, all species of the metacommunity are able to survive. Here, regional coexistence is possible because the species are spatially segregated (Fig. 4a), with each patch dominated by its best matching species.

**Fig. 5.**
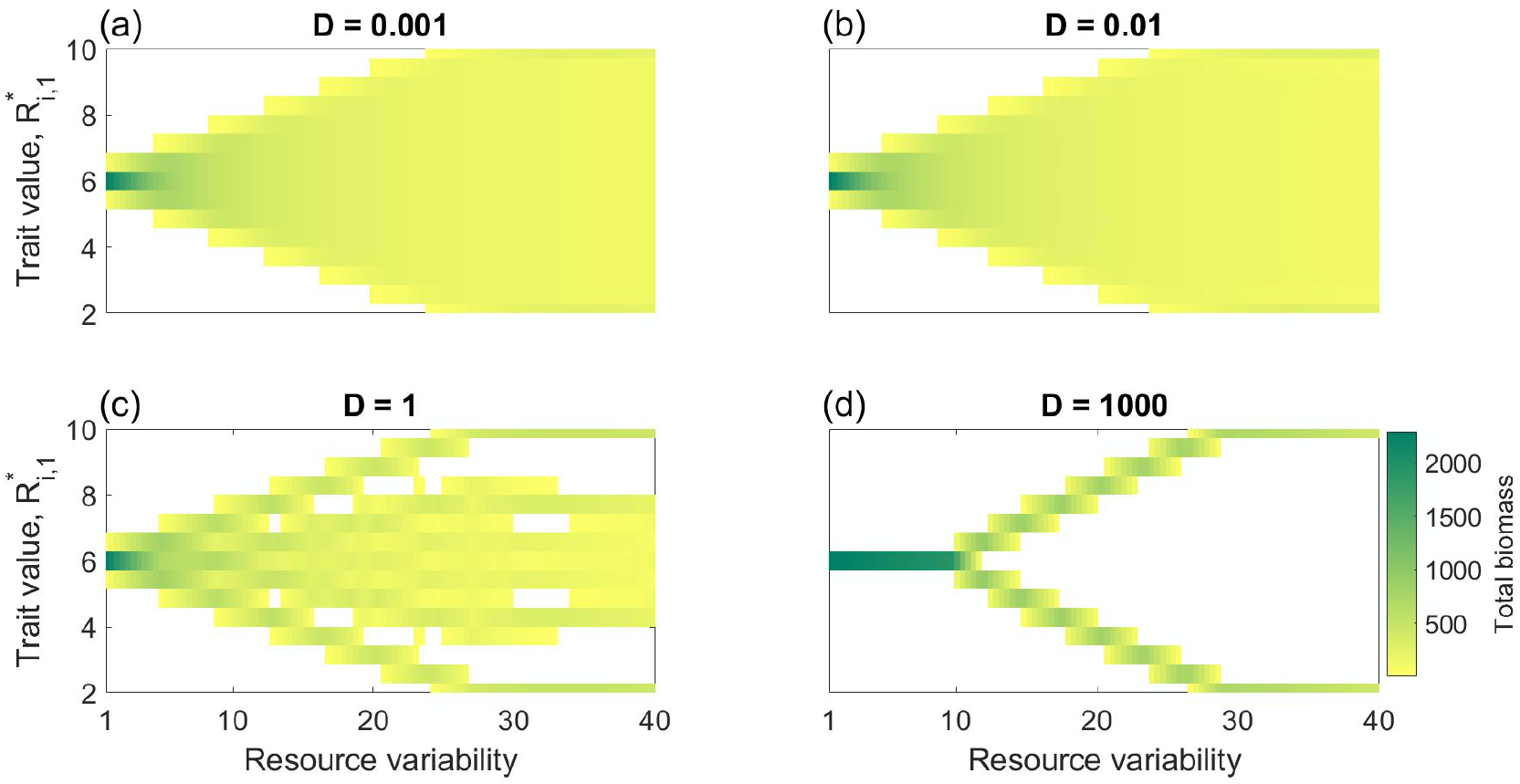
Dispersal and spatial environmental variability interact and shape competition outcomes, species richness, and community composition. The figure shows the total biomass (color coding) of a metacommunity of 15 species in equilibrium in dependence of the resource variability (Δ*S*) and the trait value (requirement, *R*\*_*i*,1_, for resource 1) for four different levels of dispersal. Parameter values as in Fig. 3.

When the dispersal rate has grown to moderate values, we observe the formation of gaps in the trait distribution, indicating the onset of trait lumping (Fig. 5c). The appearance of trait gaps is caused by increasing interspecific competition as the stronger dispersal drives species into sink locations that are already occupied by other species. With increasing dispersal this effect is amplified, further increasing the size of the trait gaps. Thereby, close to each of the two edges of the trait range a particularly large trait gap is created, separating a smaller trait range of moderate species (the ‘tongue’ from Fig. 3) and two extreme species at the ends of the trait range. Increasing dispersal even further, this large trait gap is becoming larger, successively reducing the extent of the moderate trait range. Finally, at high dispersal rates when the resource variability is sufficiently large, the moderate species are fully suppressed so that only the extreme species survive (Fig. 5d). In this range, the metacommunity is characterized by spatially homogeneous biomass distribution and the coexistence of these two extreme resource specialists. When resource variability is reduced but dispersal rate is still large, the trait difference of the two surviving species becomes smaller, until with further reduction of the resource variability a bifurcation occurs and the two surviving species merge to a single moderate species. In this range the metacommunity is characterized by a spatially homogenous distribution of this single moderate species.

### Combined effect of dispersal and resource variability on local and regional diversity

In Fig. 6 we summarize the combined effect of dispersal and resource variability on the local and regional diversity of both the small (3 species) and the large (15 species) metacommunity. At low dispersal rates, regional diversity *E*_reg_ is determined by environmental filtering (Fig. 6, top row). That is, *E*_reg_ is low, starting from *E*_reg_ = 1, when resource variability is small and it increases with increasing resource variability, approaching its maximal level defined by the size of the initial species pool. Thus, in line with the results above (Figs. 2 and 3) the largest regional diversity requires low dispersal and large environmental variability. Local diversity, however, remains small, *E*_loc_ = 1, in this range of low dispersal because in every patch the best adapted species dominates and locally outcompetes the other species.

**Fig. 6:**
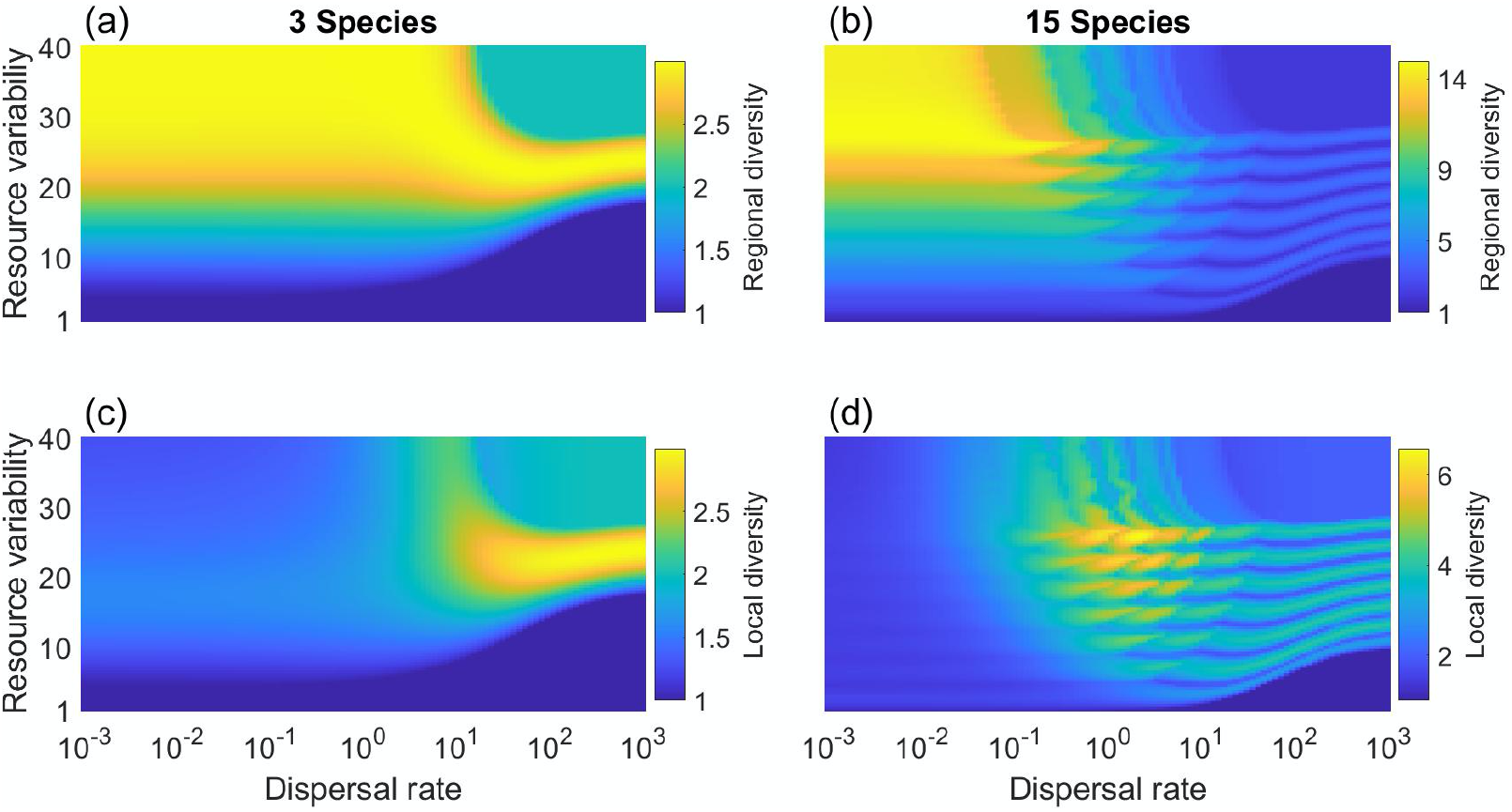
Joint effect of dispersal and environmental variability on the equilibrium diversity of metacommunity on an environmental gradient. The figure shows the regional diversity (a, b) and the local diversity (c,d) in color coding, in dependence of the dispersal rate and resource variability for a metacommunity of 3 species (left column, a,c) and 15 species (right column, b,d). Parameter values otherwise as in Fig. 3.

With increasing dispersal rate, the regional diversity in general is reduced, but the final value achieved depends on the range of resource variability. For small resource variability the regional diversity remains small, *E*_reg_ = 1, independent of the dispersal rate, as only a single species with intermediate resource requirements survives. In contrast, for very large resource variability regional diversity approaches the value *E*_reg_ = 2 for a large dispersal rate because in this limit only the two extreme species can coexist. Finally, an intermediate resource variability can create conditions balancing the competitive capabilities of the extreme and intermediate competitors, so that the regional diversity at intermediate and high dispersal rates achieves a maximum at intermediate resource variability, where both the extreme and some intermediate species coexist (see also Fig. 5c). In the system with 3 species in this situation all three species coexist, and in the system with 15 species the effective number of species rises up to 4 for large dispersal rate (see also SI Fig. 8).

The local diversity pattern deviates from the regional diversity mainly in the range of small dispersal (Fig. 6, bottom row), where local diversity is always low, because only one or two species can persist locally, while global diversity can be large in case of a large range of resource variability. With increasing dispersal rate, species distributions broaden in space and start to overlap. As a result, the local diversity increases, due to the mass effect (see Fig. 4). However, this increase is pronounced only in the range from moderate to large range of resource variability, where the regional diversity is high. When the resource variability and, therefore, the regional pool of species with positive growth in some patch is small, the mass effect becomes negligible and increasing dispersal has only a negative effect on local diversity. At high dispersal rates local diversity approaches regional diversity, as species biomasses are evenly distributed over all patches. As a consequence, at high dispersal, in a system with 3 species the local diversity is maximal at intermediate resource variability (Fig. 6c).

Note that the wave-like structure in the pattern of local and regional diversity in the right panels of Fig. 6 is no universal phenomenon, but rather a model artefact related to the discreteness of the patch model and trait distribution. The effects of this discreteness can be seen in Fig. 5d, where with a continuous increase of resource variability the dominance shifts from one species to its next neighbor on the trait axes. During these shifts the diversity measure oscillates from one (only the first species is present) to two (the first and the second species are present) and then back to one (only the second species is present). This effect also explains the small oscillations of local and regional diversity as a function of dispersal in Fig. 3.

## Discussion

Among the drivers that affect regional diversity in metacommunities two factors stand out: spatial environmental heterogeneity and dispersal. Our analysis revealed novel insights into the role of these two factors for shaping coexistence patterns, trait distributions, spatial structure and diversity along environmental gradients. In our simulations we systematically varied the variability of resource supply and dispersal rate and focused, in particular, on contrasting the response of ‘moderate’ species (with trait values in the middle of the trade-off curve) from that of ‘extreme’ species (with traits located at the two edges on the trade-off curve). We found that for small resource variability, the traits of extreme species are always filtered out by the environment and moderate species win the competition. This is because there is no matching between the traits of extreme species and the spatial environmental conditions (Cadotte and Tucker 2017). That is, the trait values of extreme species fall out of the feasible range on the spatial environmental gradient (Fig. 1). Extreme species are only the best competitors when the spatial environmental variability is large. Under such conditions, both moderate and extreme species can stably coexist at low dispersal rates. Increasing dispersal leads to an increased effect of interspecific competition, with the consequence that moderate species go extinct. This is most likely due to the fact that the traits of extreme species are located at the edges on the trait axis and, therefore, they experience competition only from one side along the gradient.

In general, the influence of dispersal depends on the level of resource variability. For small dispersal the dynamics are dominated by environmental filtering, so that in each patch the best adapted species wins. With increasing dispersal, we observe in general, a reduction in regional biodiversity, as well as the emergence of patterns in the trait distribution (trait lumping). In the limit of large dispersal for small environmental variability the winner is a moderate species with traits in middle of trade-off curve, while for large environmental variability the two extreme species located at the edges of the trade-off curve win.

Our study confirms the results of a previous analysis by Mouquet et al. (2006) who analyzed a similar model for a system of two patches and four species. In our model set-up, using a one-dimensional lattice of 50 patches and larger species pool, we were able to resolve spatial structure and finer details in the emerging trait distribution, revealing complex transitions in spatial and trait patterns with increasing dispersal. In particular, in our simulations with higher trait resolution we found remarkable structures and gaps in the trait distribution, which appeared at two different scales. On the one hand, two large trait-gaps appeared at the edges of the feasible trait range. These gaps separate the two extreme species from the ‘tongue’ of moderate species in the mid of the trait range. The size of these trait-gaps varies with the dispersal rate, leading in the limit of large dispersal to the collapse of the moderate-range trait-tongue, and thus provides a route to the coexistence state of two extreme species. On the other hand, we observed the emergence of small trait-gaps within the moderate trait-tongue. While the large trait-gaps probably are induced by boundary effects at the edges of the trait range, these small trait gaps are unlikely to be caused by the boundaries, suggesting that they may be caused by genuine pattern formation process (Meron 2015). This effect is reminiscent to effect of species lumping on a trait axis (Scheffer and van Nes 2006, Pigolotti et al. 2007) and probably is generated by similar mechanisms, as analog trait gaps arise also in alternative models of trait-based metacommunities of competing species (Doebeli and Dieckmann 2003, Leimar et al. 2008, Norberg et al. 2012).

Our simulations revealed the remarkable observation that in the limit of large dispersal and sufficiently large resource heterogeneity two resource specialists can coexist and dominate the metacommunity, despite that fact that the system in this limit effectively becomes spatially homogeneous. This is in contrast to classic resource competition theory (Tilman 1982, Chase and Leibold 2009, Koffel et al. 2016) where in a spatially homogeneous system with many species only a single best adapted consumer should survive. We remark that our results are basically just numerical observations and, in particular, for large dispersal rates we observed increasingly long transient times until the system fully equilibrated. Thus, we cannot completely rule out that our observations in the large dispersal limit are just transients. To validate our results, we have confirmed our results with various numerical ODE-solving schemes, including root-solving algorithms for equilibrium states which should be independent of transient dynamics. All these different algorithms yielded very similar outcomes, making the possibility of numerical artifacts rather unlikely. A more rigorous analysis would require analytical calculations in the limit of infinite dispersal in which spatial structure can be neglected.

Taken together, species dispersal and the spatial environmental variability strongly affect the local and regional diversity of species. Regional diversity is determined by the strength of the environmental filter while local diversity is rather determined by the competitive strength (Laliberté et al. 2014). Our results have demonstrated various routes by which dispersal affects local and regional diversity, depending also on the range of resource variability (Mouquet et al. 2006). These findings (Fig. 6) unify the two hump-shaped relationships of maximal diversity for intermediate levels of dispersal (Mouquet and Loreau 2002, 2003) and for intermediate resource variability (Kunin 1998, Mouquet and Loreau 2002, Mouquet et al. 2006).

We designed this study as a conceptual investigation. We do not aim to model a specific system and leave it as a challenge for future work to test our theoretic predictions in field data. Nevertheless, the described patterns should be relevant in a wide range of systems, as environmental gradients arise in many natural systems (Hall et al. 1992, Herbert et al. 2004, Kraft et al. 2015, Cadotte and Tucker 2017). Models of species competing for two essential resources are often applied to phytoplankton growing on mineral resources (Grover 1997, Ryabov and Blasius 2011). There, resource gradients of changing N to P resource ratios arise, for example, along estuaries or in the transition between more eutrophic coastal and more oligotrophic open ocean waters. Gradients in resource supply have also been applied to model plant distributions (Herbert et al. 2004). The role of environmental gradients for promoting biodiversity was shown in an experimental study on artificial stream ecosystems (Cardinale 2011), where a system with a higher number of available niches required a matching trait variability for an optimal use of the provided resources, while when niche opportunities were artificially removed, making all habitats uniform, the system collapsed to the dominance of a single species. Unimodal relationship between dispersal frequency and diversity were observed, for example, in a study by Kneitel and Miller (2003) who investigated communities found in the water‐filled leaves of the pitcher plant *Sarracenia purpurea*. The authors showed that dispersal among local communities can have a variety of effects on species composition and diversity at local and regional scales. In particular, increased dispersal frequencies increased regional species richness and abundance. Similar to our simulations, maximal diversity in the midrange of environmental gradients is frequently observed in the wild. This was, for example, shown in a recent study on salt-marsh plant communities along a gradient of groundwater depth and salinity from the pioneer zone to the upper saltmarsh (Bauer et al. 2021). In another experiment in stream ecosystems, Fraaije et al. (2015) investigated the interplay of dispersal versus environmental filtering on plant species distributions along riparian gradients in stream ecosystem and found a maximum diversity for intermediate values of water level.

In our conceptual study, we left out many biologically relevant aspects. We are confident that, despite these simplifications, our results are robust. Nevertheless, our study provides many avenues for model extensions and future research. Most notably, it would be interesting to investigate the influence of a variable range of trait distributions (which were here fixed to a constant range). Other possible model extensions include the addition of stochastic or seasonal influences, to extend the model to two-dimensional landscapes and heterogenous, possibly random, distributions of resource supply, and to implement different trade-offs between species resource requirements and consumption vectors. Even without extending the model, many important aspects of our model were not explored. For example, here we focused on community states in equilibrium situations, but it would be worthwhile to also study transients to equilibrium and the time scales that the system needs to reach steady state conditions in the different parameter regimes. Even though in our numerical simulations we varied among many different regimes of initial conditions, we did not systematically explore the possibility that the system may have parameter regimes with bistable behavior. Finally, it would be interesting to go beyond purely numerical investigations and to derive some of our observed patterns in analytical calculations, in particular, in the limit of large dispersal where the system simplifies as the spatial structure is eliminated.

## Acknowledgment

We thank H. Hillebrand for useful comments on the manuscript and Daniel Huber for contributing to an earlier version of this study.

This project was funded by German Research Foundation (DFG) in the Research Unit DynaCom Grant Number FOR 2716.

## Author contributions

B.B. and A.R. conceived the study and designed the model, B.B and M.M performed the numerical simulations in parallel using different programing laguages, MM prepared the results. All authors contributed to the writing.

Correspondence and requests for materials should be addressed to B.B.

## Appendix

**Fig. 7:**
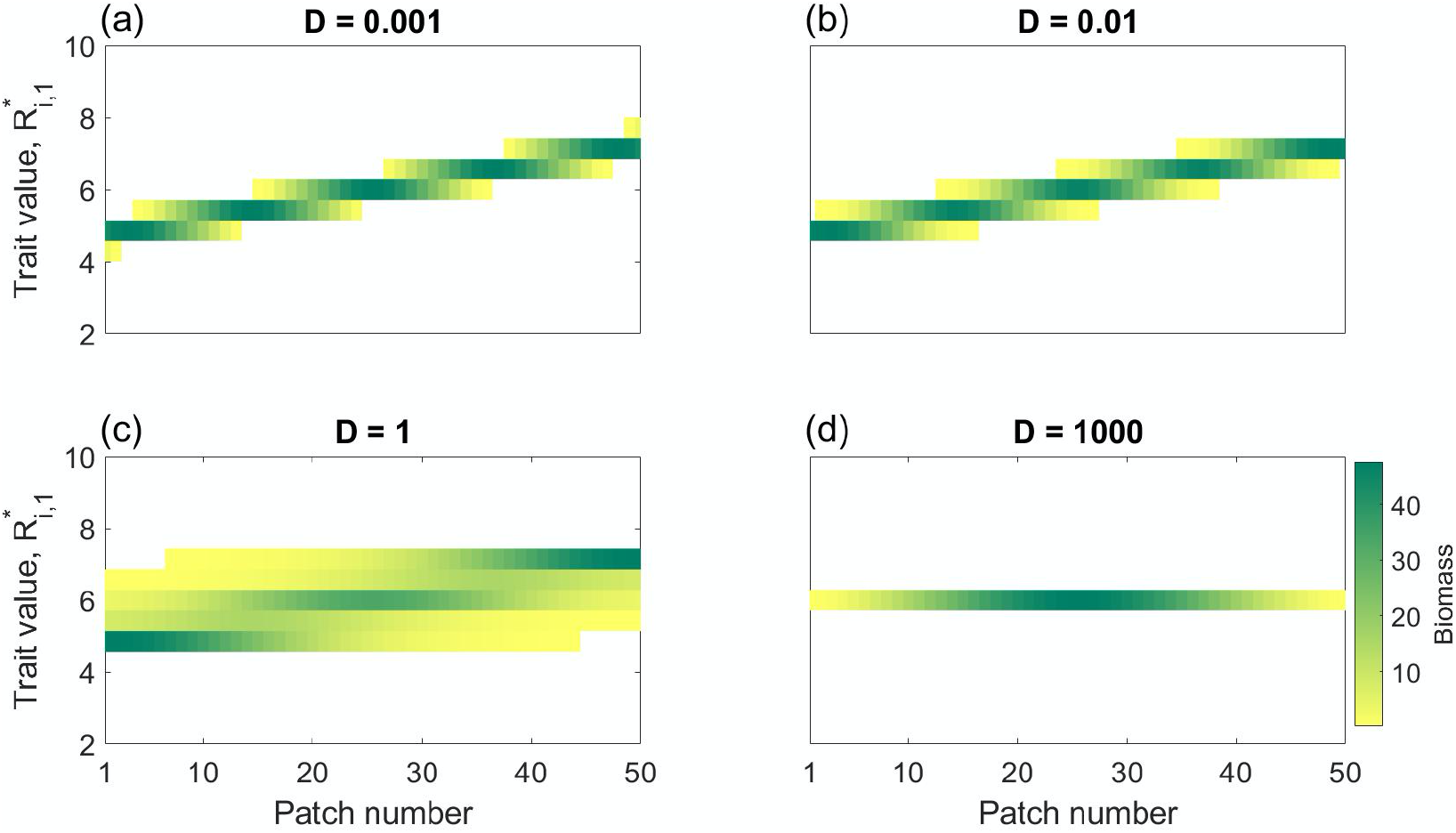
The same as Fig.4 but for a smaller resource variability of Δ*S*=8. (a-c) For small to intermediate dispersal rate the biomass distributions are similar to that of the case of a larger resource variability (Δ*S*=40, Fig. 4), with the main difference that the realized trait range is smaller. (d) For large dispersal rate (*D* = 1000), however, only a single moderate species is dominating the system.

**Fig. 8:**
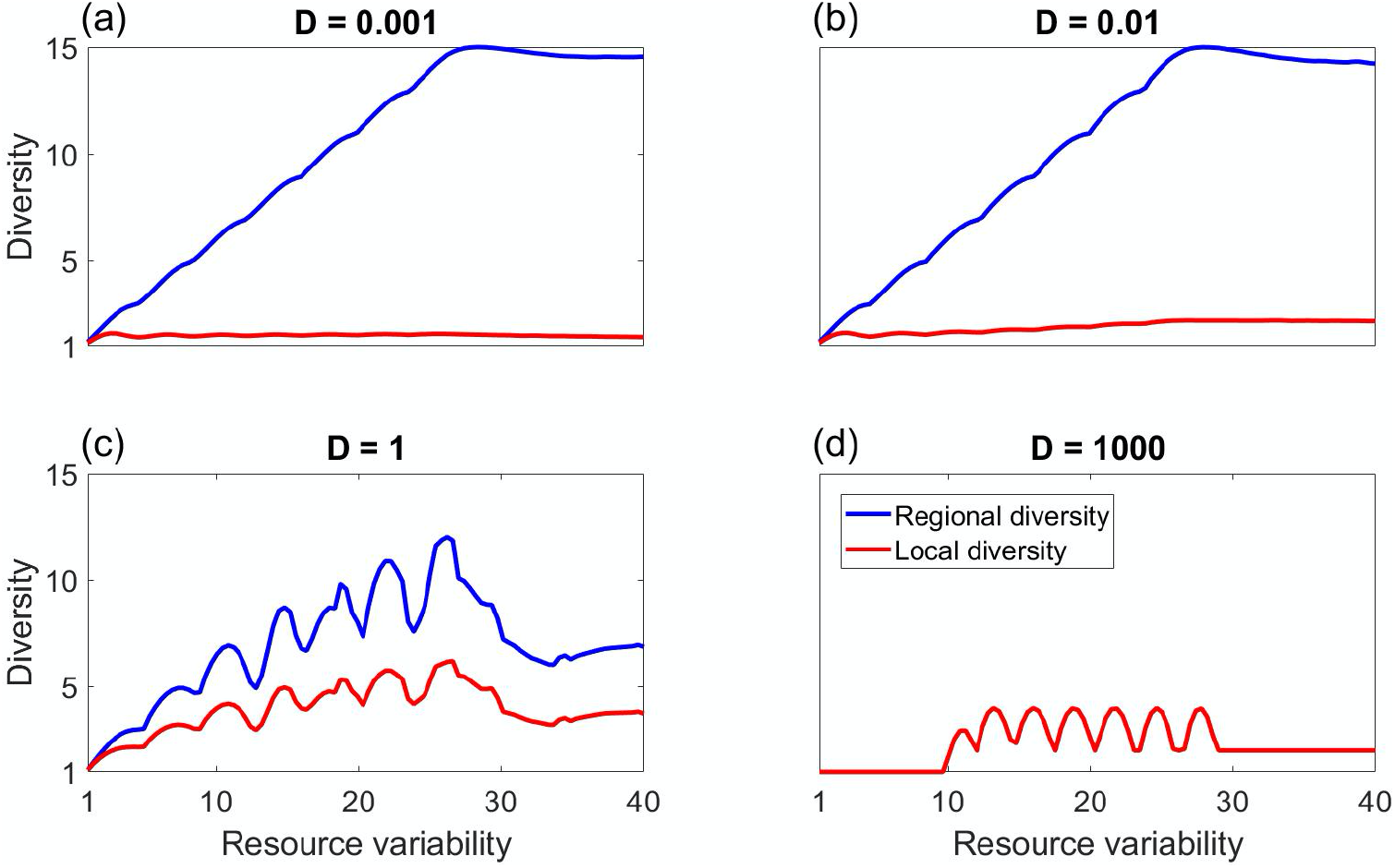
Diversity in dependence of resource variability. The figure shows the local and regional effective species number (red and blue lines) of a metacommunity of 15 species in equilibrium in dependence of the resource variability (Δ*S*) for four levels of dispersal rate. Parameter values otherwise as in Fig. 5.

## References

Abrams, P.A. and Wilson, W.G., 2004. Coexistence of competitors in metacommunities due to spatial variation in resource growth rates; does R* predict the outcome of competition? Ecology Letters, 7, 929–940.

Bauer, B., Kleyer, M., Albach, D. C., Blasius, B., Brose, U., Ferreira-Arruda, T., Feudel, U., Gerlach, G., Hof, C., Kreft, H., Kuczynski, L., Lõhmus, K., Moorthi, S., Scherber, C., Scheu, S., Zotz, G. and Hillebrand, H., 2021. Functional trait dimensions of trophic metacommunities. Ecography (accepted).

Berg, M.P., Kiers, E.T., Driessen, G., Van Der Heiden, M.A.R.C.E.L., Kooi, B.W., Kuenen, F., Liefting, M., Verhoef, H.A. and Ellers, J., 2010. Adapt or disperse: understanding species persistence in a changing world. Global Change Biology, 16, 587–598.

Bezanson, J., Edelman, A., Karpinski, S. and Shah, V. B., 2017. Julia: a fresh approach to numerical computing. SIAM Rev., 59, 65–98.

Cadotte, M.W. and Tucker, C.M., 2017. Should environmental filtering be abandoned? Trends in Ecology & Evolution, 32, 429–437.

Case, T.J. and Taper, M.L., 2000. Interspecific competition, environmental gradients, gene flow, and the coevolution of species' borders. The American Naturalist, 155, 583–605.

Cardinale, B.J., 2011. Biodiversity improves water quality through niche partitioning. Nature, 472, 86–89.

Chase, J.M. and Leibold, M.A., 2009. Ecological Niches. University of Chicago Press.

Delfau, J.B., Ollivier, H., López, C., Blasius, B. and Hernández-García, E., 2016. Pattern formation with repulsive soft-core interactions: Discrete particle dynamics and Dean-Kawasaki equation. Physical Review E, 94, 042120.

Devictor, V., Julliard, R. and Jiguet, F., 2008. Distribution of specialist and generalist species along spatial gradients of habitat disturbance and fragmentation. Oikos, 117, 507–514

Doebeli, M. and Dieckmann, U., 2003. Speciation along environmental gradients. Nature, 421, 259

Fournier, B., Mouquet, N., Leibold, M.A. and Gravel, D., 2017. An integrative framework of coexistence mechanisms in competitive metacommunities. Ecography, 40, 630–641.

Fraaije, R.G., Braak, C.J., Verduyn, B., Verhoeven, J.T. and Soons, M.B., 2015. Dispersal versus environmental filtering in a dynamic system: drivers of vegetation patterns and diversity along stream riparian gradients. Journal of Ecology, 103, 1634–1646.

Grover, J., 1997. Resource competition. Chapman & Hall. London, UK.

Haegeman, B. and Loreau, M., 2014. General relationships between consumer dispersal, resource dispersal and metacommunity diversity. Ecology Letters, 17, 175–184.

Hall, C.A., Stanford, J.A. and Hauer, F.R., 1992. The distribution and abundance of organisms as a consequence of energy balances along multiple environmental gradients. Oikos, 377–390

Herbert, D.A., Rastetter, E.B., Gough, L. and Shaver, G.R., 2004. Species diversity across nutrient gradients: an analysis of resource competition in model ecosystems. Ecosystems, 7, 296–310.

Hodapp, D., Hillebrand, H., Blasius, B. and Ryabov, A.B., 2016. Environmental and trait variability constrain community structure and the biodiversity-productivity relationship. Ecology, 97, 1463–1474.

Koffel, T., T. Daufresne, F. Massol, and C. A. Klausmeier. 2016. Geometrical envelopes: Extending graphical contemporary niche theory to communities and eco-evolutionary dynamics. Journal of Theoretical Biology, 407, 271–289.

Jost, L., 2006. Entropy and diversity. Oikos, 113, 363–375.

Kirkpatrick, M. and Barton, N.H., 1997. Evolution of a species’ range. The American Naturalist, 150, 1–23.

Kneitel, J.M. and Miller, T.E., 2003. Dispersal rates affect species composition in metacommunities of Sarracenia purpurea inquilines. The American Naturalist, 162, 165–171.

Kraft, N.J., Adler, P.B., Godoy, O., James, E.C., Fuller, S. and Levine, J.M., 2015. Community assembly, coexistence and the environmental filtering metaphor. Functional Ecology, 29, 592–599.

Kunin, W.E., 1998 Biodiversity at the edge: a test of the importance of spatial “mass effects” in the Rothamsted Park Grass experiments. Proceedings of the National Academy of Sciences, 95, 207–12.

Laliberté, E., Zemunik, G. and Turner, B.L., 2014. Environmental filtering explains variation in plant diversity along resource gradients. Science, 345, 1602–1605.

Leibold, M.A., Holyoak, M., Mouquet, N., Amarasekare, P., Chase, J.M., Hoopes, M.F., Holt, R.D., Shurin, J.B., Law, R., Tilman, D. and Loreau, M., 2004. The metacommunity concept: a framework for multi-scale community ecology. Ecology Letters, 7, 601–613.

Leibold, M.A., 2011. The metacommunity concept and its theoretical underpinnings, 163–184. In Scheiner, S.M. and Willig, M.R. eds., The Theory of Ecology. University of Chicago Press.

Leimar, O., Doebeli, M., and Dieckmann, U., 2008. Evolution of phenotypic clusters through competition and local adaptation along an environmental gradient. Evolution, 62, 807–22.

León, J. A., and D. B. Tumpson. 1975. Competition between two species for two complementary or substitutable resources. Journal of Theoretical Biology 50, 185–201.

Litchman, E. and Klausmeier, C.A., 2008. Trait-based community ecology of phytoplankton. Annual Review of Ecology, Evolution, and Systematics, 39, 615–639

Loreau, M. and de Mazancourt, C., 2013. Biodiversity and ecosystem stability: a synthesis of underlying mechanisms. Ecology Letters, 16, 106–115.

McGill, B.J., Enquist, B.J., Weiher, E. and Westoby, M., 2006. Rebuilding community ecology from functional traits. Trends in Ecology & Evolution, 21, 178–185.

Meron, E., 2015. Nonlinear physics of ecosystems. CRC Press.

Mouquet, N. and Loreau, M., 2002. Coexistence in metacommunities: the regional similarity hypothesis. The American Naturalist, 159, 420–426.

Mouquet, N. and Loreau, M., 2003. Community patterns in source-sink metacommunities. The American Naturalist, 162, 544–557

Mouquet, N., E. Miller, T., Daufresne, T. and M. Kneitel, J., 2006. Consequences of varying regional heterogeneity in source–sink metacommunities. Oikos, 113, 481–488

Norberg, J., Urban, M.C., Vellend, M., Klausmeier, C.A., Loeuille, N., 2012. Eco-evolutionary responses of biodiversity to climate change. Nature Climate Change, 2, 747–751.

Pigolotti, S., López, C. and Hernández-García, E., 2007. Species clustering in competitive Lotka-Volterra models. Physical Review Letters, 98, 258101.

Ptacnik, R., Moorthi, S.D. and Hillebrand, H., 2010. Hutchinson reversed, or why there need to be so many species. Advances in Ecological Research, 43, 1–43.

Rackauckas, C. and Nie, Q., 2017. Differentialequations.jl – a performant and feature-rich ecosystem for solving differential equations in Julia. Journal of Open Research Software, 5.

Ryabov, A.B. and Blasius, B., 2008. Population growth and persistence in a heterogeneous environment: the role of diffusion and advection. Mathematical Modelling of Natural Phenomena, 33, 42–86.

Ryabov, A.B. and Blasius, B., 2011. A graphical theory of competition on spatial resource gradients. Ecology Letters, 14, 220–228

Seabloom, E.W., Bjørnstad, O.N., Bolker, B.M. and Reichman, O.J., 2005. Spatial signature of environmental heterogeneity, dispersal, and competition in successional grasslands. Ecological Monographs, 75, 199–214

Scheffer, M. and van Nes, E.H., 2006. Self-organized similarity, the evolutionary emergence of groups of similar species. Proceedings of the National Academy of Sciences, 103, 6230–6235

Thompson, P.L., Guzman, L.M., De Meester, L., Horváth, Z., Ptacnik, R., Vanschoenwinkel, B., Viana, D.S. and Chase, J.M., 2020. A process‐based metacommunity framework linking local and regional scale community ecology. Ecology Letters, 23, 1314–1329.

Tilman, D., 1980. Resources: a graphical-mechanistic approach to competition and predation. The American Naturalist, 116, 362–393.

Tilman, D., 1982. Resource competition and community structure. Princeton University Press.

Tilman, D., 1994. Competition and biodiversity in spatially structured habitats. Ecology, 75, 2–16.

Tsakalakis, I., Blasius, B. and Ryabov, A., 2020. Resource competition and species coexistence in a two-patch metaecosystem model. Theoretical Ecology, 13, 209–221.

Fournier, B., Mouquet, N., Leibold, M.A. and Gravel, D., 2017. An integrative framework of coexistence mechanisms in competitive metacommunities. Ecography, 40, 630–41.

Wickman, J., Diehl, S., Blasius, B., Klausmeier, C. A., Ryabov, A.B., Brännström, Å 2017. Determining selection across heterogeneous landscapes: A perturbation-based method and its application to modeling evolution in space. The American Naturalist, 189, 381–95.

Wickman, J., Diehl, S. and Brännström, Å., 2019. Evolution of resource specialisation in competitive metacommunities. Ecology Letters, 22, 1746–1756.

